# Novel cytokine interactions identified during perturbed hematopoiesis

**DOI:** 10.1101/484170

**Authors:** Madison S. Krieger, Joshua M Moreau, Haiyu Zhang, May Chien, James L Zehnder, Martin A Nowak, Morgan Craig

## Abstract

Hematopoiesis is a dynamic process involving the up- and down-regulation of genes, as well as feed-back loops that stimulate or suppress circulating cytokine concentrations. More complete pictures of the gene regulatory networks that control the production of the blood system have emerged with the advent of single-cell sequencing techniques and refinements to the capabilities of immunoassays. However, information about the regulatory networks of cytokines is still lacking. A novel mathematical technique (convergent cross-mapping, or CCM) allows for the extraction of causal relationships from data, which is of crucial importance for understanding these networks. To reconstruct the cytokine networks within the hematopoietic system we measured the concentrations of 62 cytokines, platelets, and thrombopoietin from an individual with cyclic thrombocytopenia (regular oscillations in the megakaryocytes and platelets) over 84 days. Using CCM, we identified 61 previously unreported cytokine relationships. Our approach is the first broad-scale investigation into causal relationships between cytokines in the blood and suggests a new paradigm for understanding how dynamic regulation occurs during hematopoiesis.

## Introduction

The task of producing all of the body’s blood cells is a monumental one, given that some 10^11^ blood cells are derived from the hematopoietic stem cells (HSCs) every day through proliferation, differentiation, and maturation events. These processes are mediated by a cascade of cytokines responsible for amplifying or suppressing the various mechanisms that maintain basal blood cell numbers. Cytokines are a broad class of proteins that interact with blood cells at the receptor level and include stimulating and growth factors, chemoattractant chemokines, interleukins (ILs), and interferons (IFNs) that are crucial for eliciting functional hematological responses (immune-mediation, clotting, oxygen transport etc.).^1^ Different cytokines have overlapping functions to provide a measure of flexibility to the hematopoietic system to mitigate potentially dangerous cases of deficiency or receptor interference.^2^ To better understand how the hematopoietic system is maintained and responds to environmental or intrinsic stressors and/or pathological perturbations, the characterization of the relationships between these signalling proteins is necessary. Given recent advances in sequencing and computing power, the reconstruction of hematopoietic gene regulatory networks has been of increasing interest.^3–5^ The identification of the regulatory organization of the blood has enabled a better elucidation of transcriptional hematological networks,^3, 5^ and has led to the characterization of so-called ‘hubs’ (central nodes within the network graph that have many connections).^4^ However, despite these advances, obtaining a complete picture of the cytokine regulation cascades that manage blood cell counts and functions remains difficult.

Generally speaking, the feedback interactions of the cytokine/hematopoietic system paradigm function without fail throughout life. However a number of hematological pathologies are known to result from broken cytokine networks and dysfunctional blood cell production mechanisms, including the rare oscillatory diseases like cyclic neutropenia,^6^ and cyclic thrombocytopenia (CTP).^7–9^ Recently, a 53-year-old male patient with CTP (cycle period of 40 days with platelet oscillations between 1 × 10^9^/L and > 400 × 10^9^/L) was found to have a heterozygous germline mutation resulting in a loss of of c-mpl function.^10^ Previous spectral analysis identified statistically significant oscillations in platelet and neutrophil count and in and cytokine concentrations over time.^11^ Using sampling data from this patient, we endeavored to uncover the many relationships between more than 60 cytokines and platelets using standard and state-of-the-art statistical and mathematical techniques.

CTP produces oscillations of fixed periods in the megakaryocytes and platelet lines, and of TPO concentrations.^9, 12^ As we recently reported, during the manifestation of CTP, a variety of other cytokines exhibit oscillations with periods of varying length.^11^ Ordinarily, entities that co-cycle with identical periods are likely to be found to be strongly correlated. Furthermore, standard measures of correlation are symmetric in the sense that if A is strongly correlated with B, it follows that B is strongly correlated with A. However, it is often the case that A and B are strongly coupled because the dynamics of B are strongly dependent on A (or vice versa), meaning that one variable inherits the period of the other through a one-way dynamical hierarchy. This cannot be uncovered by simply measuring the correlation of the two time series. To construct a dynamical hierarchy that contains more comprehensive information than standard statistical correlation, we sought an advanced approach able to uncover one-way dynamical relationships between different cytokines. We therefore turned to convergent cross-mapping (CCM), a methodology combining dynamical systems theory and causative analysis, pioneered by Sugihara *et al.*^13–16^

Beyond simply identifying cytokine relationships, CCM also allows us to arrange cytokines in a hierarchical regulatory network. Studying properties of the network itself reveals much about how information about hematopoietic functions is communicated within the blood. Periodogram analysis then suggests the timescales of these transmissions, a further piece of information that is hard to extract with CCM alone. Our approach combining correlation analysis, periodograms, and CCM is validated by the identification of hundreds of relationships corroborated in the literature. Fascinatingly, we further uncovered a fraction of unreported relationships, substantiating our approach and suggesting worthwhile subjects for future investigation.

### Abbreviations

BDNF: brain-derived neurotrophic factor
CD40L: cluster of differentiation 40
EGF: epidermal growth factor
ENA78: C-X-C motif chemokine 5
FASL: Fas ligand
FGFB: fibrob-last growth factor-basic
GCSF: granulocyte colony-stimulating factor
GMCSF: granulocyte-macrophage colony-stimulating factor
GROA: C-X-C motif chemokine 1
HGF: human growth factor
ICAM1: intercellular adhesion molecule 1
IFNA: interferon-alpha
IL: interleukin
IL12P40: interleukin-12 subunit p40
IL12P70: interleukin-12 subunit p70
IFNB: interferon-beta
IFNG: interferon-gamma
LIF: leukemia inhibitory factor
M1P1B: macrophage inflammatory protein 1 beta
MIG: monokine induced by gamma interferon (C-X-C motif chemokine 5)
MCP: monocyte chemoattractant protein
MCSF: macrophage colony-stimulating factor
NGF: nerve growth factor
PAI1: cytokine modulation of plasminogen activator inhibitor-1
PDGFBB: platelet-derived growth factor-BB
PLTs: platelets
RANTES: regulated on activation, normal T cell expressed and secreted
SCF: stem cell factor
SDF1A: stromal cell-derived factor 1
TGFA: transforming growth factor alpha
TNF: tumour necrosis factor
TPO: thrombopoietin
TRAIL: TNF-related apoptosis-inducing ligand
VCAM1: vascular cell adhesion protein 1
VEGF: vascular endothelial growth factor

## Results

### Convergent cross-mapping reveals a robust network of interacting cytokines

The output of CCM analysis is a set of *cross-maps* (positive indicators that cytokine A has a causative effect on cytokine B) and a measure of the strength of this relationship (a real number between 0 and 1). Note that a cross-map is not a symmetric relationship; it is possible that A has a causative effect on B but that B does not have such an effect on A. Self cross-maps are considered redundant — therefore the data, which consisted of 63 objects (62 cytokines plus platelets), could yield a potential 63 × 62 = 3906 cross-maps. Our data consisted not only of cytokine levels but also measurements of the genes’ RNA transcription levels. Therefore, we further divided cross-maps into those that occur at the level of transcription (meaning A has a causative influence on both B and its RNA precursor) and those that do not. Of these 3906 possibilities, we uncovered roughly 1200 that met the standard of successfully cross-mapping two cytokines with Spearman rank-correlation on cross-mapping skill of *p* ≤ 0.05. We further filtered these results by requiring that the maximum cross-mapping skill *S* of at least one edge between the two cytokines exceeded the threshold *S*_*ϕ*→*ψ*_(24) > 0.8, where 24 is the total number of observations in the time series dataset. This left the 305 cross-maps pictured in Fig. 1. For a lengthier discussion of CCM and why the threshold of *S*_*ϕ*→*ψ*_(24) > 0.8 was used, see the **Supplementary Information**.

**Figure 1.**
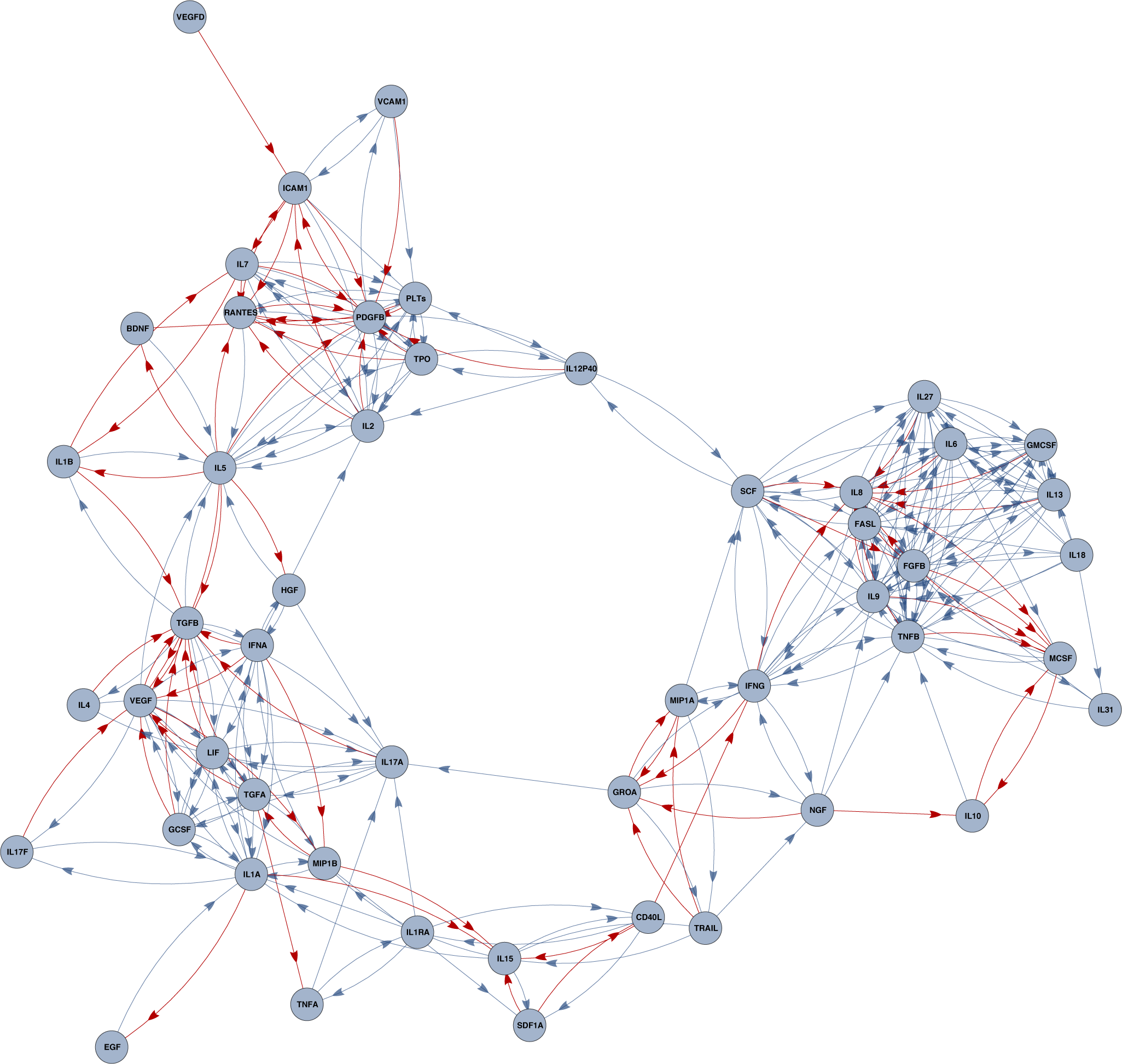
Causal relations between cytokines is revealed by convergent cross-mapping (CCM). Because positive convergent cross-mapping is not a symmetric relationship (A cross-maps B does not imply B cross-maps A), a hierarchical, directed graph can be drawn of all cytokine interactions. Here, an edge is red if the interaction is at a transcriptional level (meaning the source cytokine cross-maps the RNA levels of the target cytokine) and blue otherwise. Blue interactions could mean that the CCM signal at the transcriptional is not sufficiently strong, or that the interaction between the cytokines happens at an environmental level, for instance being inactivated by identical chemical compounds, such as inactivation by heparan sulfate in FGFB and IL8.

Analyzing the graph structure of the cytokine-cytokine interaction networks reinforces our understanding of the robustness of the immune system, even during the perturbed hematopoiesis present in this individual. At minimum, 4 cytokines (e.g. IL12P40, IL17A,CD40L and TRAIL) must be completely removed from the system before the regulatory network is meaningfully disrupted, meaning information cannot flow from one given cytokine to any other cytokine (see **Supplementary Information** for extensive analysis of the network structure provided by CCM).

While a great number of cross-maps were found at the transcriptional level, the majority are not. The absence of a transcriptional cross-map can be caused for one of three reasons: 1) because noise in the data has lowered the CCM signal of interaction at the transcriptional level below our self-imposed confidence threshold, 2) because the cytokine being cross-mapped-to (target of an edge in Fig. 1) is not produced by blood cells (such as TPO, which is produced in the liver), or 3) because the interaction between the two cytokines is genuinely at an “environmental” (rather than transcriptional) level. Examples of these latter interactions involve the competition for limited resources, such as inactivation via heparan sulfate in the case of IL8 and FGFB,^17, 18^ or an interaction between a cytokine and a cell type that produces the cross-mapped cytokine — for instance, apoptosis-inducing cytokines tend to cross-map other cytokines at this environmental level, likely because they are simply removing cytokine-producing cells from the blood.

### Elucidating the presence and timescales of between-cytokine communication

As a comparative measure, we also investigated how RNA transcription and cytokine concentrations were statistically correlated (Fig. 2). When detected, RNA expression was generally weakly correlated with cytokine concentrations in the blood, with some exception (cell surface integrins VCAM1/ICAM1 concentrations and PDGFBB gene expression, and HGF concentrations and TNFB gene expressions, for example). The general lack of information about how cytokines interact with gene expression when viewed through this lens stands in stark contrast to Fig. 1, where CCM analysis revealed a much greater amount of interaction between plasma cytokine levels and gene expression.

**Figure 2.**
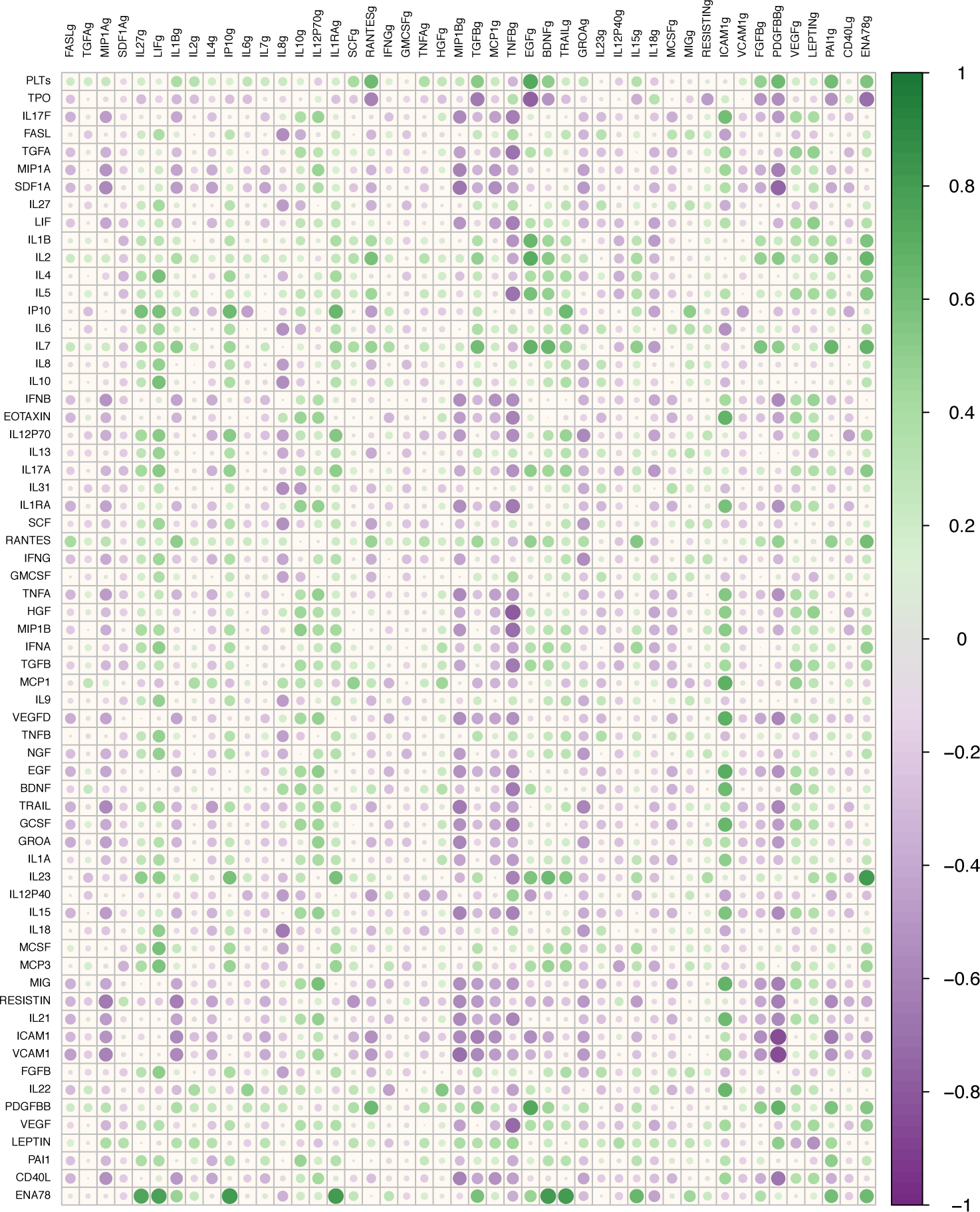
Pairwise correlation between cytokines and platelets. Pairwise Pearson correlations were calculated between platelets and cytokines (rows) and gene expressions (columns). Names appended with g indicate genes.

To compare the cross-mapping results with the pairwise Pearson correlation results, we reinterpreted the network structure of Fig. 1 as a table similar to a more usual correlation matrix, as seen in Fig. 3. Surprising differences between cytokine interactions are revealed: while certain clusters of correlated cytokines are found via convergent cross mapping to also have a causative effect on one another, other correlated clusters are almost entirely absent in the CCM picture. This could either be because these cytokines indeed have no direct causative effect on one another, and co-cycle due to forcing by some other cue, or because noise in the data has reduced the cross-mapping strength of these relationships between our imposed threshold. We found that many uncorrelated cytokines are actually seen to cross-map. Given that these relationships are virtually invisible under the standard umbrella of statistical techniques, this finding further emphasizes the advantages gained by the application of the CCM method.

**Figure 3.**
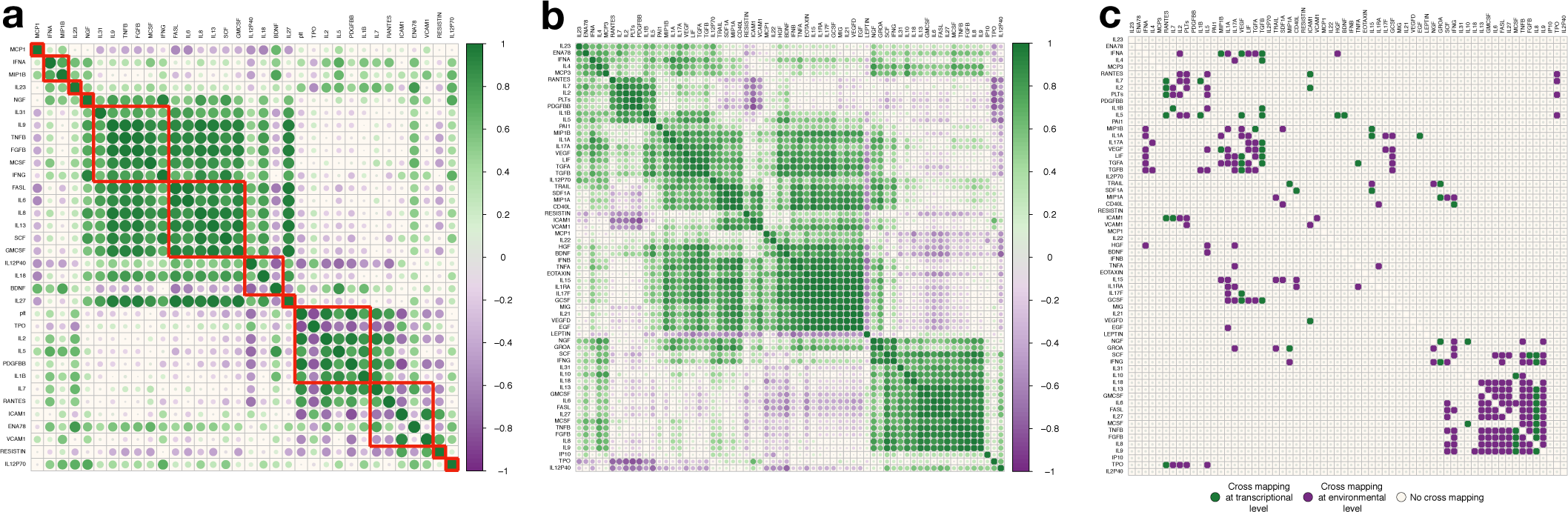
Statistical and causal relationships revealed through Pearson correlation and convergent cross-mapping. **a)** Pairwise correlation results for cycling cytokines and platelets, ordered according to period class. From top to bottom (and equivalently, from left to right), each row(column) is ordered according to the length of their cycling period. Red squares classify individual period classes (entities cycling with the same periods). Deep purple indicates a perfectly negative correlation, deep green indicated perfectly positive correlation. **b)** Pairwise Pearson correlations were calculated between each cytokine (and platelets). Gradient scale as in a. **c)** Alternative formulation for Figure 1. Since the structure is not symmetric (A crossmaps B does not imply B crossmaps A), a mark indicates that the cytokine along the vertical axis (horizontal cytokine names) has a causative influence on the cytokine along the horizontal axis (vertical cytokine names). If this influence has a strong CCM signature at the transcriptional level, the mark is green. If the CCM signature is seen at the population/environmental level but not at the transcriptional level, the mark is purple.

### Cytokine clusters emerge according to both correlation similarity and the period of oscillations

We have previously identified eight distinct cycling periods within the data (hereafter called “period classes”).^11^ Restricting our analyses to those cytokines (and platelets) that exhibited these statistically significant oscillations, and grouping them according to period class (denoted by the red squares in Figure 3a), we looked for strong positive (in phase) or negative (out of phase) correlations within each period class. The distinction of in or out of phase can be made by combining the spectral Lomb-Scargle analysis^11^ and the correlation results of Figure 3b. We observed that the vast majority of correlations within each period class are positive, indicating that most cytokines within the same class are oscillating in phase, with the exception of the associations between BDNF/IL12P40, ICAM1/IL7, and ICAM1/RANTES, and *all* molecules within the 38.9565-day period class and TPO.

The strength of the calculated correlation coefficients between any two entities is denoted by both the intensity of the colour (green or purple) and the size of the circle in Figure 3b.^19^ Unsurprisingly, platelets and TPO are perfectly negatively correlated (i.e. TPO concentrations fall as platelet numbers increase and vice versa). The cell surface integrins ICAM1 and VCAM1, and RESISTIN display strong negative correlation with the RANTES/IL7/IL2/platelet/PDGFBB subgroup only while most other cytokines are positively pairwise correlated. However when a statistically significant relationship exists, leptin, IP10, TPO and IL12P40 are almost entirely negatively correlated with every other cytokine. While the full analysis revealed correlations between RANTES and IL7 in the second cluster, the period class cross-mapping identified causal relationships between RANTES, IL7, and VCAM1/ICAM1, which were found to cluster in the fourth correlation group.

### Exploiting period classes to expose novel causal relationship structures

Using our previous Lomb-Scargle results,^11^ we explored the cross-maps between cytokines within period classes, displayed in Fig. 4. Given the degree of pairwise correlations within each class, we anticipated the causal networks to be complete graphs (each node connected to every other by a unique edge). The and 31.87 day classes almost achieve completeness, save for one or two edges (in the 29.22 class, for example, IFNG and IL31/MCSF do not cross-map, nor do MCSF and IL31, whereas SCF does not cross-map to IL13 in the 31.87 class), however interesting structures emerge in the 35.06, 38.96 (the period of oscillations in the platelets and TPO), and 43.86 classes where graphs were not complete. In the 38.96 day class, IL1B only sends and receives information from IL5 and functions as a controller outside of the complete subgraph formed by IL5, IL2, TPO, PDGFB, and circulating platelet concentrations. Similarly, in the 43.83 class, VCAM1 sits atop the complete subgraph comprised of ICAM1, IL7, and RANTES, and interacts solely with VCAM1. Most curiously, there are no edges between the members of the 35.06 class (IL18, IL12P40, BDNF), despite them being statistically correlated (see Fig. 3a).

**Figure 4.**
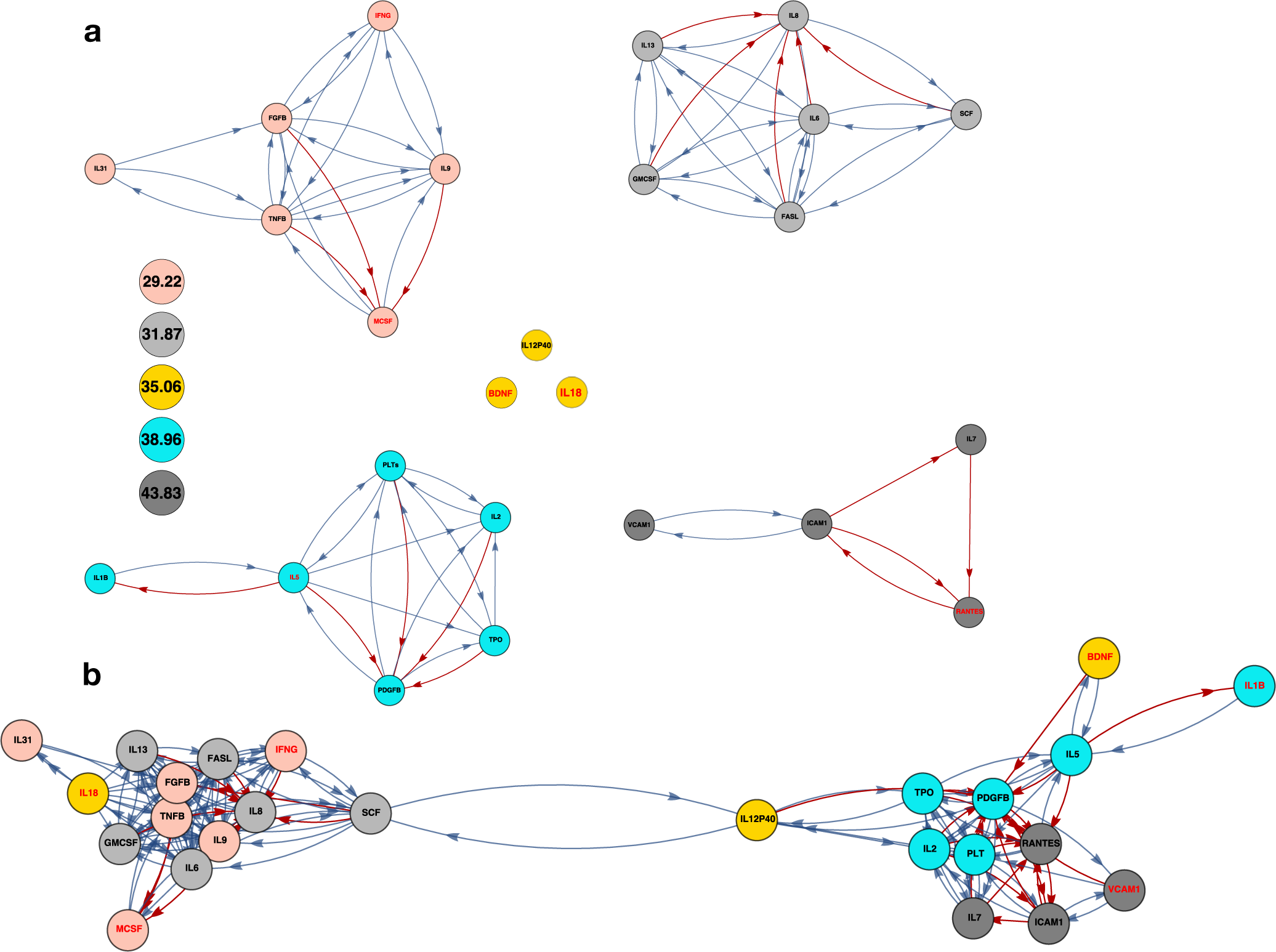
Causal relationships amongst period class pairs and cycling cytokines. **a)** Combining the periodogram analysis and convergent cross-mapping analyses reveals a causal network among elements of cytokine classes that have identical periods. Cytokines are represented in the network as nodes, with node colour assigned according to period (legend bottom-left). Nodes with black text labels cycle with their assigned period with *α* = 0.05. Nodes with red text labels cycle with their assigned period with *α* = 0.5. Directed edges indicate convergent cross-mapping: an edge directed from node A to node B indicates that, at optimal embedding dimension and library size, the cross-mapping skill *S* (see **Supplementary Information**) indicating the information contained about the state of cytokine A in the state of cytokine B is greater than 0.8. In this panel, information about causality is restricted to elements of the same periodogram class. **b)** Convergent cross-mapping analysis also uncovers links between cytokines that belong to different period classes. Here edges are coloured with the same colour as their source node. When this information is included, a hierarchical network is created between cytokines that cycle at different rates. Red arrows indicate that RNA expression is also cross-mapped between nodes.

### Establishing novel cytokine relationships through CCM analysis

Of the 305 cross-maps identified using the *S*_*ϕ*→*ψ*_(24) > 0.8 as a minimal threshold, we identified 60 previously unknown relationships between cytokines (see **Supplemental Data** for a complete list of cross-maps and identification methodology, as well as citations from the literature for the known relationships uncovered by CCM). Thus 19.6% of the cross-maps identified through convergent cross-mapping with truncation at *S*_*ϕ*→*ψ*_(24) > 0.8 have not previously been reported in the literature. These novel interactions are listed in Table 1 and include a variety of interleukins, and growth and necrosis factors. Although CCM analysis easily differentiates between scenarios in which a given cytokine A governs B and C without B and C interacting and those in which B directly interact with C, it is unable to differentiate the causal chain *A* → *B* → *C* from *A* → *C* with surety (see **Supporting Information**). We therefore include the minimum number of relationships that are already known (in the literature) to distinguish the proposed novelty. As the number of intervening known edges becomes larger, it becomes less plausible that this edge is a false positive, and therefore these are the relationships that merit future exploration the most.

**Table 1.** Previously unreported relationships revealed through convergent cross-mapping. Of the ∼1200 causal relationships identified through convergent cross-mapping (see **Supplementary Information** for details), we identified 305 relationships by truncating the maximal cross-mapping skill with the minimal threshold of *S*_*ϕ*→*ψ*_(24) > 0.8. In addition, we have indicated the number of intervening cytokines along known pathways: as this number increases, the likelihood that the novel relation is an artifact of CCM analysis decreases. Pairs highlighted in light grey were identified by both CCM and direct Pearson correlation (with *ρ* > 0.8, see **Supplementary Information**). Of the 305 causal associations found by CCM, 60 are previously unreported (19.6%). The best candidates for future investigation are highlighted in dark grey/white text. Highlighted in medium grey are those identified by both methods that are also strong candidates for further study. A more thorough exploration of these novelties is provided in the **Supplementary Material**.

| Cytokine 1 | Cytokine 2 | # intervening | Cytokine 1 | Cytokine 2 | # intervening |
| --- | --- | --- | --- | --- | --- |
| BDNF | PDGFB | 1 | IL27 | TNFB | 1 |
| FASL | FGFB | 1 | IL31 | FGFB | 1 |
| FASL | IL9 | 1 | **IL31** | **TNFB** | **2** |
| FASL | SCF | 1 | MCSF | IL9 | 1 |
| FGFB | FASL | 1 | MIP1A | SCF | 1 |
| FGFB | IFNG | 1 | MIP1A | TRAIL | 1 |
| FGFB | IL13 | 1 | MIP1B | SDF1A | 1 |
| FGFB | IL27 | 1 | MIP1B | TGFA | 1 |
| FGFB | IL9 | 1 | **PDGFB** | **ICAM1** | **2** |
| GROA | IFNG | 1 | **PDGFB** | **IL12P40** | **2** |
| ICAM1 | PDGFB | 1 | PDGFB | VCAM1 | 1 |
| IFNA | TGFA | 1 | **PLTs** | **IL5** | **2** |
| IFNG | FGFB | 1 | PLTs | PDGFB | 1 |
| IFNG | GROA | 1 | SCF | FASL | 1 |
| IFNG | IL9 | 1 | SCF | TNFB | 1 |
| **IL9** | **FASL** | **2** | SDF1A | IL15 | 1 |
| IL9 | FGFB | 1 | TGFA | IFNA | 1 |
| IL9 | IFNG | 1 | **TGFA** | **IL17A** | **1** |
| IL9 | MCSF | 1 | TGFA | IL1A | 1 |
| IL9 | TNFB | 1 | TGFA | MIP1B | 1 |
| IL12P40 | PDGFB | 1 | TGFA | TNFA | 1 |
| **IL12P40** | **PLTs** | **2** | **TGFA** | **VEGF** | **2** |
| IL12P40 | TPO | 1 | **TNFB** | **IL27** | **2** |
| IL13 | FGFB | 1 | TNFB | IL31 | 1 |
| IL15 | SDF1A | 1 | TNFB | IL9 | 1 |
| IL17A | TGFA | 1 | **TNFB** | **SCF** | **2** |
| IL17A | IL1A | 1 | TPO | IL12P40 | 1 |
| IL18 | FGFB | 1 | TRAIL | MIP1A | 1 |
| IL18 | TNFB | 1 | **VCAM1** | **PDGFB** | **2** |
| **IL27** | **FGFB** | **1** | **VCAM1** | **PLTs** | **2** |
|  |  |  | VEGF | TGFA | 1 |

Viewed through a different lens, we may say that CCM is 80% *accurate* — that is, it indicates causal interactions which have been independently corroborated by previous research 80% of the time. To gauge whether CCM is really an effective method to untangle interactions from time-series data, we compared this measure of accuracy to direct Pearson correlations (interpreting pairs with *ρ* > 0.8 to represent true interactions, just as we require the CCM predictive strength at maximum library to be larger than 0.8), and also to the null constructed by drawing pairs of cytokines uniformly-at-random and then checking against the literature. In the null case, we selected 100 cytokine-cytokine pairs at random and compared them to the previous reports, finding that 37% percent of them were accurate, i.e., were corroborated by at least one experimental paper. The number of pairs with Pearson correlation *ρ* > 0.8 was 278, similar to the 305 found by CCM with the same cutoff criterion; however, only 58% of these were corroborated by the literature. By this metric, then, CCM does seem to offer a substantial increase in interaction-inference accuracy over traditional methods. Further, CCM offers information as to the directionality of reported interactions, as opposed to Pearson correlation, which is undirected by definition. Most of the novelties suggested by CCM occur twice, with one arrow in each direction. If the results of CCM are instead treated as an undirected graph, then 19 duplicates are removed, leaving 41 unconfirmed connections, and the accuracy increases to 87%.

## Discussion

Buoyed by improvements to sequencing techniques and computational power, we are now beginning to understand how an intricate regulatory network of interacting cytokines and gene transcription in various cell lineages controls hematopoiesis. At present, a great deal of attention in this regard is paid to genetic regulation at the intracellular level. However a multitude of cytokines also transmit information as they coordinate both the production and function of the body’s blood cells. Cytokine and blood cell concentrations fluctuate daily, but additional information about the dynamic nature of the hematopoietic system is fully revealed during pathological conditions. By leveraging information obtained during dynamical disease, we sought to establish the structure of the causal cytokine relationships that present in this patient with perturbed platelet homeostasis.

Our approach uncovered many previously corroborated cytokines relationships, which validates the methodology we employed. Most causal connections were found to be environmental rather than transcriptional, a conclusion mirrored in the standard Pearson correlation measurements that are much sparser than the full pairwise correlation matrix between cytokines. In a previous analysis, we identified 12 period classes of cytokines cycling with the same duration. Combining these classes with our correlation results allowed us to identify cytokines cycling in- and out-of-phase in a given class. However, the causal communication between cytokines only comes into focus when we constructed the cross-maps within each class. We established that the networks of each class were not complete, and identified “master” transmitters of information within most classes. Most notably, we revealed ∼1200 total successful cross-maps in the full dataset, with 305 cross-maps remaining after truncation at the *S*_*ϕ*→*ψ*_(24) > 0.8 level of significance. Of these, we identified 61 previously unreported relationships that are strong candidates for future investigation. However, it would be unfair to say that the CCM analysis is somehow superior to the analysis based on statistical correlations. For example, when we plot the results of Fig. 1 in a table side-by-side with the correlation tables in Fig. 3, it is clear that the output of CCM analysis is much sparser. Further, on its own, CCM contains no information about the period of cycling of a given cytokine or co-cycling between cytokines, nor does it indicate whether a cross-mapping from A to B indicates up-regulation or down-regulation of B due to A.

Notwithstanding these limitations, we were able to broadly assess causal cytokine relationships in the hematopoietic system, despite studying only a single person, given the rich sampling and highly dynamical nature of this individual’s disease. Future work should incorporate both healthy individuals and a wider sample of people with similar dynamic pathologies, including cyclic neutropenia and chronic myeloid leukemia. This study nonetheless presents a new and powerful way to study the cytokinetic (and genetic) connections within the blood, and suggests that there is much more to discover than can be found using standard statistical means.

## Methods

### Cell counts and cytokine assays

Blood samples were drawn every 3-4 days for 84 days total (24 samples total over two cycles) from a 53-year old individual with a previously-described presentation of cyclic thrombocytopenia related to a heterozygous germline mutation resulting in the loss of c-mpl function. Blood transcriptome profiles were performed on whole blood using 3SEQ (3’-end RNA sequencing for expression quantification). Platelet poor plasma was isolated to measure thrombopoietin concentration via ELISA and a panel of 62 cytokines using a Luminex immunoassay.

### Determining causal relationships using convergent cross-mapping

For each of the measured 62 cytokines and their gene expression data, as well as platelets and thrombopoietin, we computed the cross-mapping to all other platelets/cytokines applied Sugihara *et al.’s*^13^ convergent cross-mapping (CCM) approach. CCM was developed to determine *causal* relationships in time series without reference to the usual statistical correlation.^16^ A detailed description of the mathematical foundations of convergent cross-mapping and a description of the algorithm are provided in the **Supplementary Information**. For each of the 63 × 62 = 3906 (self-maps are redundant) total possible cross-maps, we calculated the cross-mapping skill as a function of library length *L* (which cannot exceed 24, the total number of measurements) and embedding dimension *E* according to the algorithm described in the **Supplementary Information**. We then performed Spearman rank-correlation on cross-mapping skill and *L* for each cross-mapping to isolate those that showed convergent cross-mapping (by keeping only those with *p* ≤ 0.05). This subset of cross-maps was then examined by eye to remove cases where the Spearman correlation assigned a *p*-value less than 0.05, but the cross-mapping skill was not sufficiently monotonic in *L* to ensure accurate forecasting at differing library lengths.

### Correlations between measures

To understand the statistical associations between platelets, thrombopoietin, and the 62 cytokines and their genes, we quantified their pairwise Pearson correlations using the **cor** function in R^20^ and plotted the results with **corrplot**.^19^ Hierarchical clustering according to dissimilarity was applied to group statistically similar cytokines.

## Acknowledgements

MC is funded by a Discovery Grant from the Natural Sciences and Engineering Research Council (NSERC) of Canada and was previously supported by an NSERC Postdoctoral Fellowship. MC and MSK were supported by grant DP5OD019851 from the Office of the Director at the National Institutes of Health to their PI. JMM is grateful to the Human Frontiers Science Program Long-Term Fellowship for funding. We thank Jeff Gerold, Sim Sinai, Alison Hill, Michael Mackey, Tony Humphries — for helpful comments and suggestions.

## Contributions

MSK conceived and designed the CCM study, performed the CCM analysis and wrote the manuscript; JMM analyzed the complete network graph; HZ conceived and designed the sampling, cell counts, and cytokine assays, carried out the experimental work, and wrote the manuscript; MChien conceived and designed the sampling, cell counts, and cytokine assays, carried out the experimental work, and wrote the manuscript; JLZ and MAN oversaw and wrote the manuscript; MCraig conceived and designed the CCM study, performed the statistical analyses, and wrote the manuscript. All authors critically reviewed each version of the manuscript and approved the final version for submission.

## Conflict-of-interest disclosure

All authors have no conflicts of interest.

### Correspondence

Morgan Craig, Département de mathématiques et de statistique, Université de Montréal, Pavillon André-Aisenstadt C.P. 6128, succursale Centre-ville, Montréal (Québec) H3C 3J7 Canada

