## Supplemental Information for "Novel cytokine interactions identified during perturbed hematopoiesis"

### Supplementary Information: Novel cytokine interaction networks identified during perturbed hematopoiesis

#### Convergent cross-mapping

Granger causality[1] is an idea emerging from economics as a means to extract information from two time series beyond their (instantaneous) correlation. The original method, which was unrelated to attractors of dynamical systems, involved comparing the power spectra of two time series and constructing expressions for the *causal strength* and *causal lag* of variable A and variable B based on their respective spectra. Here the invocation of "causality" is related to the ability to forecast the state of B, given the state of A. The causal strength of A on B represents the correlation between a prediction of the state of B, only given information about A, and the actual value of B at the time point being forecasted. This is therefore a non-symmetric relationship: A may have causal strength over B while at the same time B has no causal strength over A. The causal lag represents the difference between the point in the time series of A where a forecast is being constructed for B, and the actual time point in the B time series where this forecast has the greatest causal strength. The idea of convergent cross-mapping is to extend Granger's original construction to a new context: that of variables that are explicitly coupled in an underlying system of nonlinear differential equations with deterministic components.

If some part of the time trajectory of a system is the flow of a deterministic equation, a useful object of study is a manifold towards which the majority of initial conditions will evolve asymptotically in time (the system's attractor). Understanding the attracting manifold $\mathbf{M}$ requires that the attractor be properly embedded in some higher-dimensional space $\mathbf{N}$. If the embedding is improper (either because the derivative of the embedding map $F : \mathbf{M} \to \mathbf{N}$ is not injective or not smooth, or fails to be a topological embedding), information about "nearness" of trajectories over time will be inaccurate and forecasting the future behaviour of the system based on the current position on the attractor will fail. Convergent cross-mapping relies on two theorems central to the embedding of attractors: Whitney's theorem[2] and Takens' theorem.[3] Whitney's theorem guarantees that the maximal dimension of $\mathbf{N}$ for which a proper embedding of $\mathbf{M}$ will exist is $\dim(\mathbf{N}) \leqslant 2\,\dim(\mathbf{M})$. To properly embed the attractor, one must find a space of proper embedding dimension in which each orthogonal subspace contains observations of the attractor, $\mathbf{M}$. To properly resolve the attractor, in some cases, $\mathbf{M}$ is a sufficient number of "obvious" observation functions, whether spatial or otherwise. The insight of Takens' theorem is to take delays of a trajectory on the attractor as independent observation functions. Given a single observation function $\phi(t)$ for the attractor that obeys certain generic properties, an embedding can be constructed using $\nu \equiv \dim(\mathbf{N})$ lags of (generic) length $\tau$, creating a $\nu$-dimensional vector at each time point $t$ in the trajectory: $(\phi(t-\tau), \phi(t-2\tau), ...\phi(t-\nu\tau))$.

The principle of convergent cross-mapping arises from the consideration of two observation functions $\phi(t)$, $\psi(t)$ that are somehow connected by an underlying nonlinear dynamical system. This connection is such that some information about the attractor of $\phi(t)$ is contained in the attractor of $\psi(t)$ (or vice-versa), or both observation functions contain some (possibly unequal) amount of information about each other. Hence the concept of Granger causality in the context of attractor cross-mapping — if the attractor underlying $\phi(t)$ contains some information about the attractor underlying $\psi(t)$, thereby reflecting coupled dynamics in some nonlinear system, then to remove the information $\phi(t)$ from the Universe would inherently diminish our ability to forecast the future of $\psi(t)$. The ability of $\phi(t)$ to accurately forecast $\psi(t)$ given the number of observations $L$ for both observation functions is here given by a function called the cross-mapping skill $S$ of $\phi$ to $\psi$:

$$S_{\phi\to\psi}(L) \in [-1, 1], \tag{S1}$$

where generically speaking $S_{\phi\to\psi} \neq S_{\psi\to\phi}$. This function is constructed as follows.

1. The attractors of $\phi(t)$ and $\psi(t)$ are both reconstructed according to Takens' theorem with embedding dimension $E$. In principle, $E$ can be as large as $L$, the number of observations; the optimal embedding dimension is defined simply as the value of $E$ for which the value of the cross-mapping skill function $S$ is maximized.

2. For each time point $T = 1, 2, \ldots, L - E$ in the delay-reconstructed manifold from $\phi(t)$, the nearest $E + 1$ neighbours in the $\mathbf{L}_2$ norm of the point corresponding to time $T$ in the high-dimensional manifold are recorded. $E + 1$ is the smallest requisite number of neighbours to ensure a generic $E$-dimensional simplex.

3. For a given point $\phi_E(T)$, each neighbour in this $E + 1$ neighbour-set is assigned a weight that exponentially decreases based on its distance from $\phi_E(T)$.

4. These neighbours (which are each an integer between 1 and $L$, their index in the time series $\phi(t)$) are now used to "predict" the value $\psi(T)$. The predicted value, called $\psi^*(T)$, is simply the sum of the value of $\psi(t)$ at the neighbours' indices weighted by the weights they were assigned when they were indices for neighbours of $\phi(T)$.

5. The cross-mapping skill is taken to be the Pearson correlation between the predicted value of $\psi(T)$ based on $\phi(T)$ and the actual value: $S_{\phi\to\psi}(L) = \rho(\psi^*(T), \psi(T))$.

The cross-mapping is said to be convergent if the cross-mapping skill increases with increasing number of available data points, $L$, indicating that the predictive power of $\phi(t)$ on $\psi(t)$ is increasing with increasing data thus confirming that some information about the attractor of $\phi(t)$ is reflected in the attractor of $\psi(t)$.

#### Network properties of causal relations determined by convergent cross-mapping

Here we include several relevant measures of the network induced on cytokines by convergent cross-mapping. Note that in the theory of convergent cross-mapping, a cross-mapping with skill less than a certain threshold is not necessarily insignificant, as the strength of an interaction could be obscured by noise in measurement. The critical observation is the convergence with increasing amounts of available data, $L$. We impose our cutoffs arbitrarily to manage the network size, both for the purpose of visualization and computation, and also to keep only the results that tend to have the highest confidence.

We depict several important measures here graphically, as well as tabulating the cytokines with the highest scores by certain metrics. In Fig. S1 we plot the in-degree and out-degree distributions. Other critical graph-theoretic properties are tabulated in Table S1. We paid particular interest to betweenness-centrality,[4] highlighting cytokines that are likely to receive and then transmit information and may therefore represent important mediators in the cytokine hierarchy. The strongest nodes by this measure are also tabulated in Table S2 and mainly including interferons, growth factors, and cytokines related to necrosis. Other metrics examined included out-closeness-centrality[5] (Table S3), HITS authority[6] (Table S4), and HITS hub-ranking[6] (Table S5). All of these metrics attempt to gauge the importance of a node for transmitting information to others, and are therefore based on the distribution of edges between the node of interest and its neighbors, next-neighbors, etc. Each measure differs from the others based on the feature of interest — for instance, between-closeness may be said to focus on the role of a node as a mediator of information (receiving and then forwarding information from other nodes) while out-closeness might be said to focus on the role of a node as a generator or forwarder of information only. It is therefore noteworthy that some of the most important cytokines by these differing metrics are the same — for instance, VEGF appears as one of the top ten cytokines by all metrics employed.

| Property | Value |
|---|---|
| Clustering coefficient | 0.541 |
| Average path length | 3.437 |
| Diameter | 7 |
| Density | 0.124 |
| Modularity | 0.590 |

**Table S1.** Additional graph properties of the cross-mapping network at cutoff strength $S > 0.8$.

| Cytokine | Scaled between-centrality |
|---|---|
| IFNG | 1.0000 |
| IL18 | 0.9843 |
| IFNA | 0.8405 |
| SCF | 0.7728 |
| CD40L | 0.7401 |
| IL5 | 0.6989 |
| VEGF | 0.5770 |
| IL12P40 | 0.5047 |
| TRAIL | 0.4883 |
| NGF | 0.4860 |

**Table S2.** Between-centrality ranking in the $S > 0.8$ network.

| Cytokine | Scaled out-closeness-centrality |
|---|---|
| CD40L | 1.0000 |
| IL17F | 0.9918 |
| GROA | 0.9758 |
| MIP1A | 0.9528 |
| VEGF | 0.9528 |
| TRAIL | 0.9453 |
| SDF1A | 0.9380 |
| IL15 | 0.9380 |
| IL1RA | 0.9308 |
| TGFB | 0.9237 |

**Table S3.** Out-closeness-centrality ranking in the $S > 0.8$ network.

| Cytokine | Scaled authority |
|---|---|
| IFNA | 1.0000 |
| VEGF | 0.9038 |
| IL17F | 0.8573 |
| EGF | 0.8149 |
| HGF | 0.8041 |
| IL1A | 0.7983 |
| IL17A | 0.7971 |
| TNFA | 0.7864 |
| IL15 | 0.7368 |
| MIP1B | 0.7260 |

**Table S4.** HITS authority ranking in the $S > 0.8$ network.

| Cytokine | Scaled hub factor |
|---|---|
| IL1RA | 1.0000 |
| MIP1B | 0.9611 |
| LIF | 0.9565 |
| VEGF | 0.9373 |
| IL1A | 0.9248 |
| TNFA | 0.9094 |
| IL15 | 0.8947 |
| GCSF | 0.8768 |
| TGFA | 0.8628 |
| TGFB | 0.7518 |

**Table S5.** HITS hub ranking in the $S > 0.8$ network.

#### Comparison with the literature

In the main text, we provided a rationale for considering CCM to be at least, in part, an advance over traditional techniques for inferring interactions — whereas older approaches, such as simply measuring the Pearson correlation coefficient, give roughly 50% to 60% "accuracy" (meaning 50% to 60% of its inferred relationships can be substantiated from other experiments), CCM in this case has given nearly 80%-90% accuracy by the same metric. Here we begin to agglomerate references to the literature to support our claim. The references to pairs identified by CCM that might be less well-known in the literature are collected in Table S6.

| Cytokine 1 | Cytokine 2 | Reference | Cytokine 1 | Cytokine 2 | Reference |
|---|---|---|---|---|---|
| MIP1B | VEGF | [7] | FASL | GMCSF | [8] |
| TGFA | GCSF | [9] | TGFA | IL1A | [10] |
| IL8 | FGFB | [11, 12] | TGFA | LIF | [13] |
| BDNF | IL5 | [14] | TPO | IL7 | [15] |
| SCF | IL8 | [16] | IFNG | IL8 | [17] |
| MCSF | FGFB | [18] | IL7 | PDGFB | [19] |
| RANTES | PDGFB | [20] | SDF1A | CD40L | [21] |
| IL18 | IL31 | [22] | IL12P40 | SCF | [23] |
| IL2 | RANTES | [24] | IL7 | RANTES | [25] |
| MIP1A | IFNG | [26] | NGF | IFNG | [27] |
| LIF | IFNA | [28] | MIP1B | IFNA | [29] |
| NGF | IL9 | [30] | IL15 | TRAIL | [31] |
| IL17f | IL1A | [32] | MIP1B | IL15 | [33] |
| IL6 | FGFB | [34] | SCF | FGFB | [35] |
| GCSF | LIF | [36] | TPO | IL2 | [37] |
| ICAM1 | IL2 | [38] | SCF | IL6 | [39] |
| FGFB | IL6 | [40] | IL7 | PLTs | [41] |
| FASL | IFNG | [42] | FASL | IL27 | [43] |
| ICAM1 | PLTs | [44] | IL18 | FASL | [45] |
| IL1A | IL15 | [46] | IL1A | IL17F | [47] |
| IL5 | TPO | [48] | RANTES | PLTs | [49] |

**Table S6. Known relationships in Fig. S2 from the literature.** Fig. S2 illustrates Fig. 1 with edges re-colored according to their status in the literature. When we considered a relationship to be less well-known, we sought out corroboration in the literature, provided in the citations here.

#### Difficulty for CCM in identifying causation in series versus in parallel

CCM, as a novel tool in causative analysis, is an exciting and promising avenue to extract networks of interactions from data on dynamical systems. However, it is possible for it to mis-identify relationships in certain settings. It has been demonstrated that CCM can correctly identify settings in which two variables are both strongly forced by a third, external variable.[50–53] However, it is possible for CCM to erroneously identify relationships as in parallel when in fact they are in serial, mitigated by other variables (see Fig. S3) — in fact, this is a common issue in causal inference, no matter the technique.[54–56] Since a large number of our "novel" cytokine interactions in Table 1 could easily be understood instead as strong forcings via only one intervening cytokine, along known pathways, it is very important to keep this fact in mind.

#### The network induced by Pearson correlation and its accuracy

In the previous sections, we established the network of interactions found by CCM, assessed its accuracy and discussed possible mechanisms by which a number of the "novel" interactions it found could in fact be false positives. However, we still maintain that it is a major improvement upon alternative techniques. In order to support that claim, we constructed a similar network by simply keeping all cytokine pairs whose Pearson correlation exceeded 0.8 (in analogy with the CCM network in which we required the predictive strength to exceed 0.8). That network is plotted in Figure S4. Whereas roughly 10%-20% of CCM's edges were unsubstantiated by the literature, 40%-50% of the Pearson network's edges are unsubstantiated. Those that were found have citations in Table S7.

| Cytokines | Reference | Cytokines | Ref | Cytokines | Ref | Cytokines | Ref |
|---|---|---|---|---|---|---|---|
| IL17F-LIF | [57] | IL17F-IFNB | [58] | IL17F-TNFA | [59] | IL17F-MIP1B | [60] |
| IL17F-TGFB | [61] | IL17F-GCSF | [62] | IL17F-IL15 | [63] | IL17F-MIG | [64] |
| IL17F-IL21 | [65] | FASL-IL31 | [66] | TGFA-IL1RA | [67] | TGFA-TNFA | [67] |
| TGFA-HGF | [68] | TGFA-TGFB | † | TGFA-EGF | [69] | TGFA-BDNF | [70] |
| TGFA-IL15 | [71] | MIP1A-IFNB | [72] | MIP1A-IL1RA | [73] | MIP1A-TNFA | [74] |
| MIP1A-GCSF | [75] | MIP1A-IL15 | [76] | MIP1A-MIG | [77] | MIP1A-IL21 | [78] |
| MIP1A-CD40L | [79] | SDF1A-ICAM1 | † | IL27-IL10 | [80] | LIF-TNFA | [81] |
| LIF-HGF | [82] | LIF-IL15 | [83] | LIF-CD40L | [84] | IL1B-IL2 | † |
| IL4-IFNA | [85] | IL4-NGF | [86] | IL6-IL9 | [87] | IL8-IL10 | [88] |
| IL8-NGF | [89] | IL8-IL18 | [90] | IFNB-IL1RA | [91] | IFNB-HGF | [92] |
| IFNB-EGF | [93] | IFNB-TRAIL | [94] | IFNB-MIG | [95] | IFNB-IL21 | [96] |
| EOTAXIN-TNFA | [97] | EOTAXIN-GCSF | [98] | EOTAXIN-IL15 | [99] | EOTAXIN-IL21 | [100] |
| IL13-SCF | [101] | IL1RA-HGF | [102] | IL1RA-EGF | [103] | IL1RA-BDNF | [104] |
| IL1RA-TRAIL | [105] | IL1RA-GCSF | [106] | IL1RA-IL15 | [107] | IL1RA-VEGF | [108] |
| SCF-IL18 | [109] | IFNG-NGF | [110] | TNFA-TGFB | [111] | TNFA-TRAIL | † |
| TNFA-GCSF | [112] | TNFA-MIG | [113] | TNFA-IL21 | [114] | TNFA-VEGF | [115] |
| TNFA-CD40L | [116] | HGF-TGFB | [117] | HGF-GCSF | [118] | HGF-VEGF | [119] |
| MIP1B-TGFB | [120] | TGFB-BDNF | [121] | TGFB-IL15 | [122] | VEGFD-VCAM1 | [123] |
| VEGFD-CD40L | [124] | EGF-GCSF | [125] | EGF-IL15 | [126] | BDNF-GCSF | [127] |
| TRAIL-IL21 | [128] | TRAIL-CD40L | [129] | GCSF-IL21 | [130] | GCSF-CD40L | [131] |
| IL15-MIG | [132] | IL15-IL21 | [133] | IL15-VCAM1 | [134] | MIG-IL21 | [135] |
| MIG-CD40L | [136] | IL21-VCAM1 | [137] | VCAM1-CD40L | [138] | | |

**Table S7. Citations for pairs identified by Pearson correlation** In order to compare the accuracy of the graphs induced by keeping pairs with CCM strength greater than 0.8 and Pearson correlation greater than 0.8, we had to construct both networks and see what fraction of edges in each were corroborated by external experiments. This table contains the citations found for less commonly-known pairs picked up by direct Pearson correlation.

#### The null hypothesis for relationships between cytokines

To define a baseline "accuracy" to compare Pearson correlations and CCM, we selected 100 cytokine pairs uniformly-at-random (ignoring duplicates and pairs that were a cytokine interacting with itself) from all possible pairs, and sought out experimental literature confirming that they had an explicit effect on one another (in either direction). The pairs chosen, and references to the literature when they were corroborated, can be found in Table S8.

| **Cytokines** | **Ref** | **Cytokines** | **Ref** | **Cytokines** | **Ref** | **Cytokines** | **Ref** |
|---|---|---|---|---|---|---|---|
| RESISTIN-TRAIL | X | FASL-SCF | X | TGFB-RANTES | [139] | TRAIL-EGF | X |
| PLTs-PAI1 | [140] | IL2-SDF1A | X | IL31-IL5 | X | HGF-IL13 | [141] |
| VEGF-IL22 | [142] | FASL-IL12P40 | [143] | IL8-RANTES | X | MCP1-VEGF | [144] |
| RANTES-PAI1 | X | IFNB-IL1A | X | TRAIL-TGFB | [145] | GROA-TRAIL | † |
| IL4-LIF | [146] | CD40L-IL2 | [147] | IL17F-IL9 | [148] | VCAM1-GCSF | [149] |
| IL4-IL31 | [150] | TNFA-PAI1 | [151] | IL1B-IL21 | [152] | BDNF-IL31 | X |
| IL5-RANTES | † | HGF-IL27 | X | TNFB-IL6 | † | IL1RA-VEGFD | X |
| IL7-IFNB | X | NGF-IL8 | [89] | MIG-PAI1 | X | IL12P40-IL13 | X |
| VEGFD-SDF1A | X | IL22-VCAM1 | X | MIG-IL8 | X | TNFB-TPO | X |
| RANTES-EOTAXIN | X | IL2-IL4 | X | RANTES-MIP1A | X | RANTES-BDNF | [153] |
| GROA-IL1A | X | MIP1A-IL8 | X | MCP3-TPO | X | PDGFBB-IL12P40 | X |
| HGF-VEGFD | [154] | IL6-GMCSF | † | LEPTIN-MCP1 | X | IFNA-IL6 | X |
| HGF-EGF | X | GCSF-IL1RA | [106] | IL31-CD40L | [155] | IL17F-VCAM1 | X |
| FGFB-IL13 | † | TPO-IL12P40 | X | NGF-ICAM1 | [156] | TRAIL-IFNA | [157] |
| PAI1-BDNF | [158] | IFNG-MIG | [159] | IL2-IL1RA | X | MIP1A-LIF | X |
| TGFB-TRAIL | [160] | TGFB-IL12P70 | [161] | IL1B-IL10 | [162] | GMCSF-VCAM1 | X |
| IFNA-VEGF | [163] | GCSF-PLTs | [164] | TNFB-IL1RA | X | IL22-PLTs | [165] |
| GROA-TGFA | X | IL4-IL23 | [166] | TNFA-IL1B | [167] | IL13-NGF | [168] |
| BDNF-IL17F | X | TGFA-IL12P70 | X | SCF-MIG | X | IL23-PAI1 | X |
| SDF1A-NGF | X | MCSF-RANTES | X | MIP1A-IL8 | X | BDNF-LEPTIN | X |
| IL7-GCSF | X | IFNB-GCSF | X | IL13-MCSF | X | TNFA-RANTES | X |
| TNFA-IL18 | X | IL12P70-MCSF | X | IL5-IL12P40 | X | IL7-IL22 | X |
| IL4-IFNB | X | MIP1A-VEGFD | X | GROA-MCP1 | X | SDF1A-TRAIL | X |
| IL13-GCSF | X | LIF-IL23 | X | MIP1B-MCSF | X | IL23-FGFB | X |
| FGFB-NGF | X | MCP3-VEGFD | X | IL4-IL7 | X | HGF-IL2 | X |

**Table S8. Citations for pairs identified by the random null** In order to compare the accuracy of the graphs induced by keeping pairs with CCM strength greater than 0.8 and Pearson correlation greater than 0.8, we also had to compare this with a null hypothesis in which pairs were selected completely at random. These are those pairs and all substantiating literature references, when they can be found (a mark of X indicates no such literature was found, while a mark of † indicates that this relationship was considered to be commonly known in the immunology literature). 37 of the randomly chosen pairs had some basis in fact.

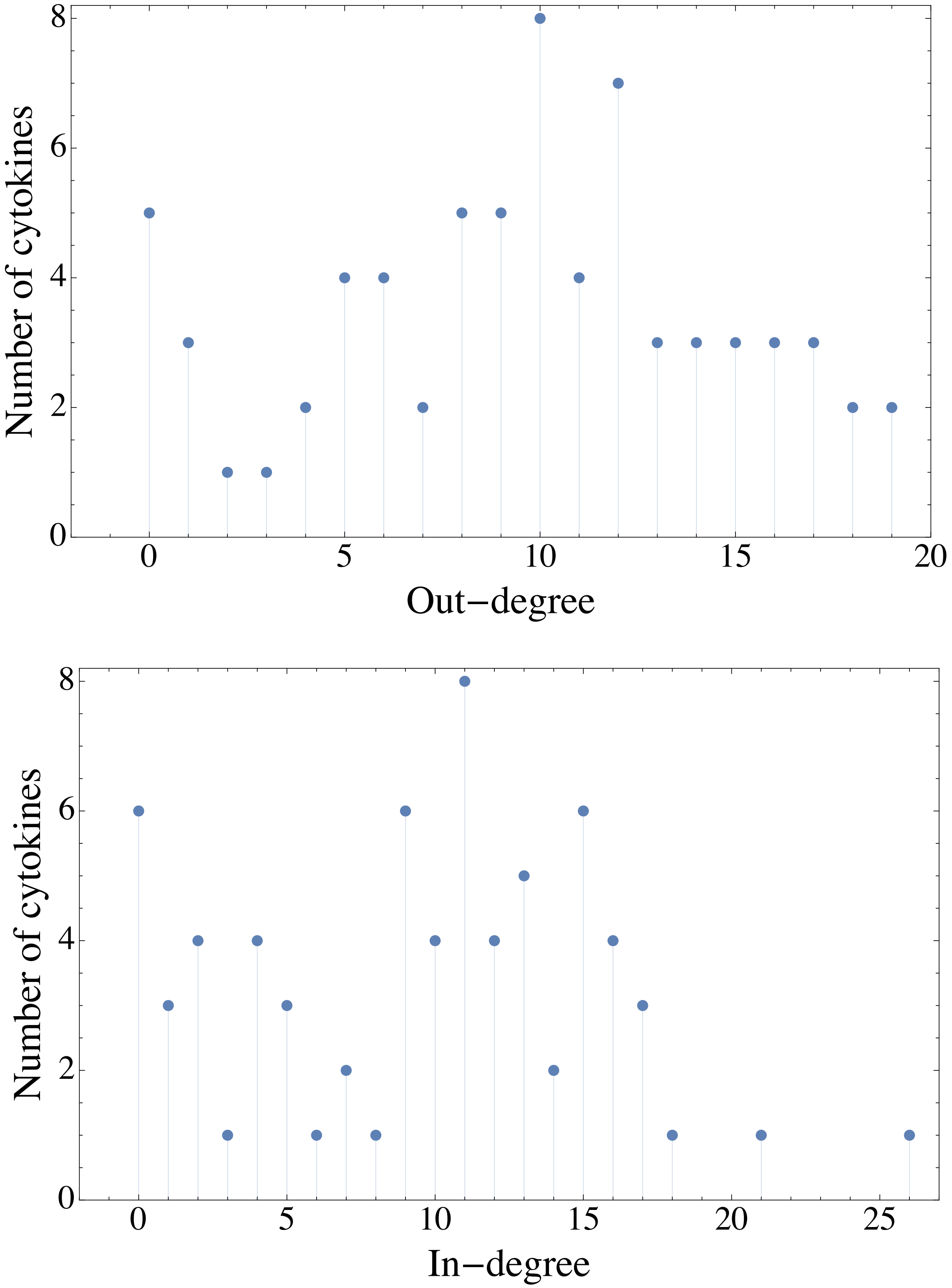


**Figure S1. Degree distributions for cytokines in network induced by convergent cross-mapping.** A larger network of cytokines induced by convergent cross-mapping which is still statistically meaningful, but difficult to visualize, is that in which all edges with maximum skill above 0.8 are maintained. Degree distributions are shown for this network. No meaningful power law is observed in the distribution.

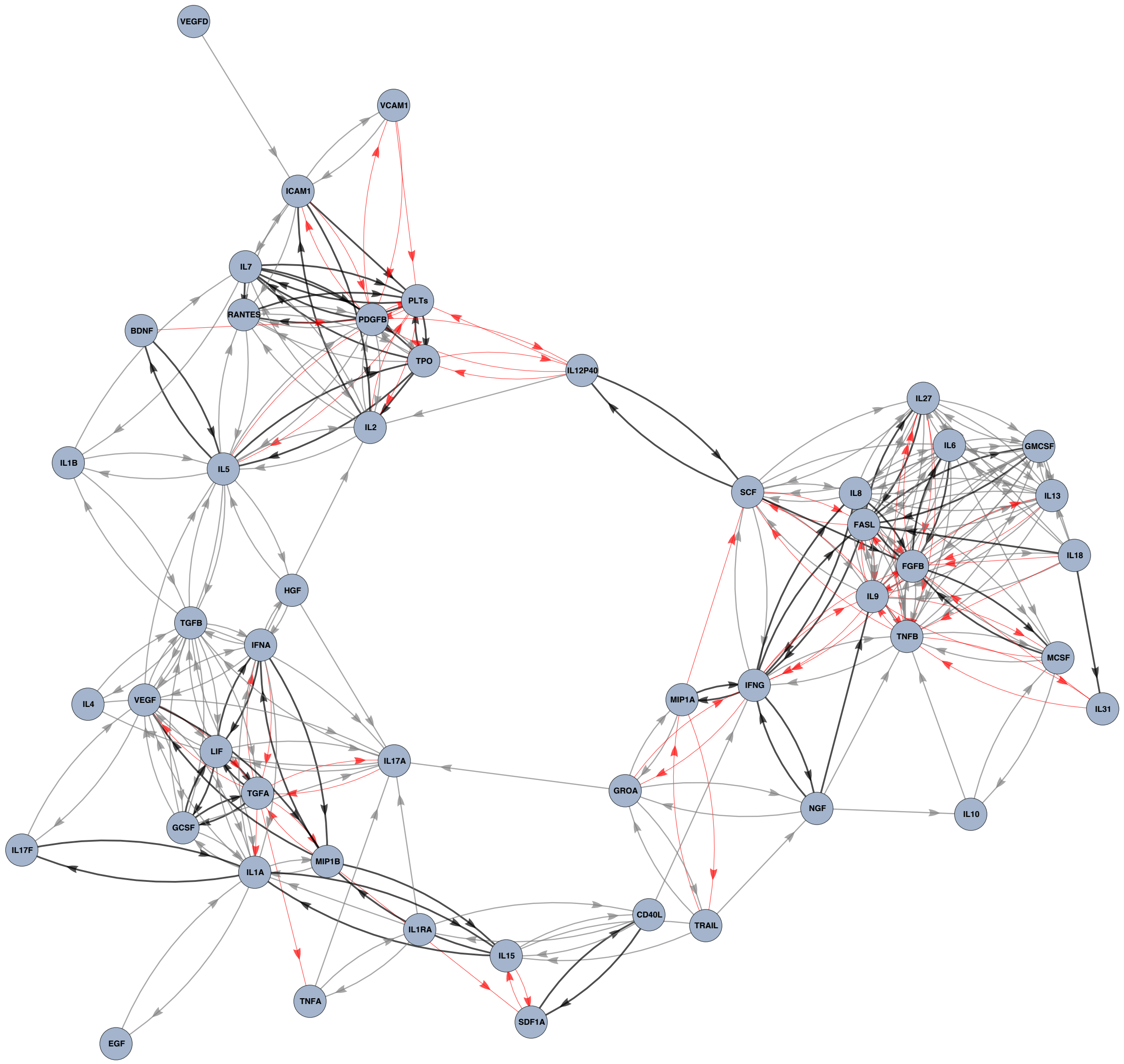


**Figure S2. What is known in the literature from Fig. 1.** This is the equivalent of Figure 1, but with edges re-colored according to their status in the literature. Gray edges indicate relationships that we considered standard, that is, the interaction is so well-known in the immunology field that no citation is needed. Black edges are supported by citations given in Table S6. Red edges indicate pairs identified by CCM for which we could find no corroborating experiments in the literature.

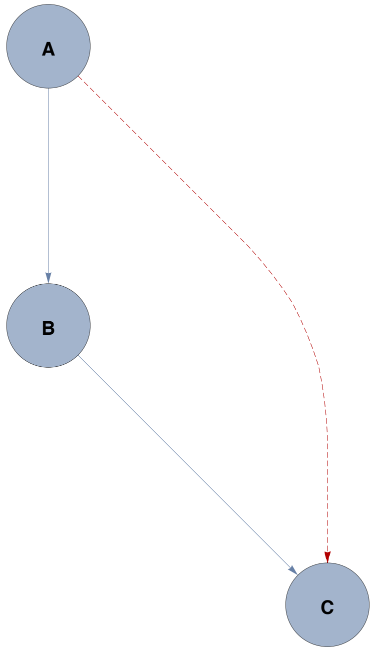


**Figure S3. An example of spurious links identified by CCM** Depending on the relative strength of interactions, CCM will either correctly ignore this edge or spuriously indicate it with positive cross-mapping.

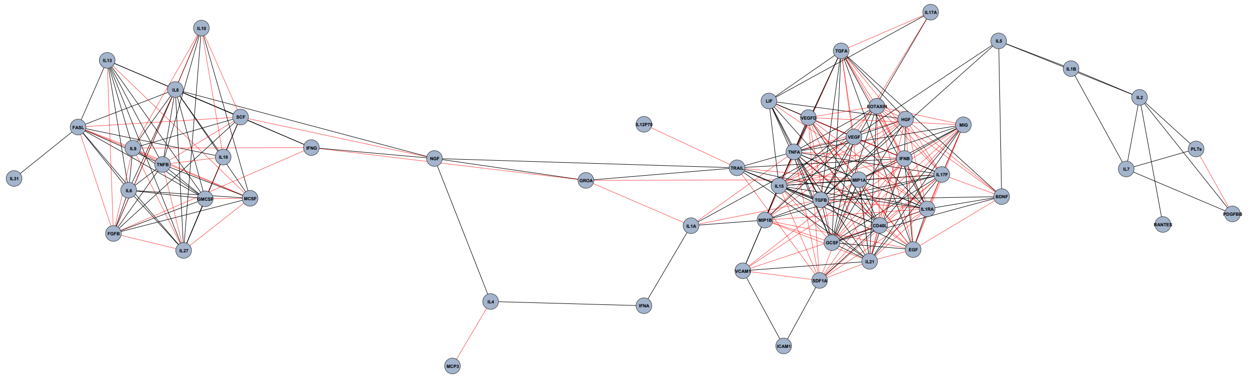

**Figure S4. The equivalent of Figure 1 when correlation is employed** Instead of drawing edges between cytokines when their CCM predictive strength at maximum library exceeds 0.8, we could do the equivalent process where an (undirected) edge is drawn between two cytokines if their Pearson correlation coefficient exceeds 0.8. While both procedures create graphs with similar numbers of edges, 57% of these edges are represented in the literature, versus the 87% of undirected edges found via CCM. Here, black edges represent known interactions from the literature, listed in Tables S6 and S7. Red edges indicate pairs for which we could find no corroborating evidence in the literature.

74. C. D. L. Ramos, C. Canetti, J. T. Souto, J. S. Silva, C. M. Hogaboam, S. H. Ferreira, and F. Q. Cunha, "Mip-1[ccl3] acting on the ccr1 receptor mediates neutrophil migration in immune inflammation via sequential release of tnf- and ltb4," *Journal of Leukocyte Biology*, vol. 78, no. 1, pp. 167–177.

75. T. Queto, Z. F. M. Vasconcelos, R. A. Luz, C. Anselmo, A. A. A. Guiné, P. M. R. e. Silva, J. Farache, J. M. T. Cunha, A. C. Bonomo, M. I. C. Gaspar-Elsas, and P. Xavier-Elsas, "G-csf suppresses allergic pulmonary inflammation, downmodulating cytokine, chemokine and eosinophil production," *Life Sciences*, vol. 88, no. 19, pp. 830–838, 2011.

76. E. M. Bluman, K. J. Bartynski, B. R. Avalos, and M. A. Caligiuri, "Human natural killer cells produce abundant macrophage inflammatory protein-1 alpha in response to monocyte-derived cytokines," *J Clin Invest*, vol. 97, pp. 2722–2727, Jun 1996.

77. M. Thapa, R. S. Welner, R. Pelayo, and D. J. J. Carr, "Cxcl9 and cxcl10 expression are critical for control of genital herpes simplex virus type 2 infection through mobilization of hsv-specific ctl and nk cells to the nervous system," *J Immunol*, vol. 180, pp. 1098–1106, Jan 2008.

78. M. Croce, V. Rigo, and S. Ferrini, "Il-21: a pleiotropic cytokine with potential applications in oncology," *J Immunol Res*, vol. 2015, pp. 696578–696578, 2015.

79. Y. Guo, A. M. Walsh, U. Fearon, M. D. Smith, M. D. Wechalekar, X. Yin, S. Cole, C. Orr, T. McGarry, M. Canavan, S. Kelly, T.-A. Lin, X. Liu, S. M. Proudman, D. J. Veale, C. Pitzalis, and S. Nagpal, "Cd40l-dependent pathway is active at various stages of rheumatoid arthritis disease progression," *The Journal of Immunology*, 2017.

80. G. Murugaiyan, A. Mittal, R. Lopez-Diego, L. M. Maier, D. E. Anderson, and H. L. Weiner, "Il-27 is a key regulator of il-10 and il-17 production by human cd4+ t cells," *J Immunol*, vol. 183, pp. 2435–2443, Aug 2009.

81. C. S. Levy, V. Slomiansky, A. Gattelli, K. Nahmod, F. Pelisch, M. Blaustein, A. Srebrow, O. A. Coso, and E. C. Kordon, "Tumor necrosis factor alpha induces lif expression through erk1/2 activation in mammary epithelial cells," *Journal of Cellular Biochemistry*, vol. 110, no. 4, pp. 857–865.

106. M. Schwabe, A.-M. Hartert, H. Bertz, and J. Finke, "Treatment with granulocyte colony-stimulating factor increases interleukin-1 receptor antagonist levels during engraftment following allogeneic stem-cell transplantation," *European Journal of Clinical Investigation*, vol. 34, no. 11, pp. 759–765.

114. V. Phan-Lai, Y. Dang, E. Gad, J. Childs, and M. L. Disis, "The antitumor efficacy of il2/il21-cultured polyfunctional neu-specific t cells is tnf/il17 dependent," *Clinical Cancer Research*, 2015.

121. A. Sometani, H. Kataoka, A. Nitta, H. Fukumitsu, H. Nomoto, and S. Furukawa, "Transforming growth factor-1 enhances expression of brain-derived neurotrophic factor and its receptor, trkb, in neurons cultured from rat cerebral cortex," *Journal of Neuroscience Research*, vol. 66, no. 3, pp. 369–376.

122. M. Benahmed, B. Meresse, B. Arnulf, U. Barbe, J.-J. Mention, V. Verkarre, M. Allez, C. Cellier, O. Hermine, and N. Cerf-Bensussan, "Inhibition of tgf-β signaling by il-15: A new role for il-15 in the loss of immune homeostasis in celiac disease," *Gastroenterology*, vol. 132, pp. 994–1008, Mar 2007.

123. T. Duong, K. Koltowska, C. Pichol-Thievend, L. Le Guen, F. Fontaine, K. A. Smith, V. Truong, R. Skoczylas, S. A. Stacker, M. G. Achen, P. Koopman, B. M. Hogan, and M. Francois, "Vegfd regulates blood vascular development by modulating sox18 activity," *Blood*, vol. 123, no. 7, pp. 1102–1112, 2014.

124. S. Kiriakidis, E. Andreakos, C. Monaco, B. Foxwell, M. Feldmann, and E. Paleolog, "Vegf expression in human macrophages is nf-b-dependent: studies using adenoviruses expressing the endogenous nf-b inhibitor ib and a kinase-defective form of the ib kinase 2," *Journal of Cell Science*, 2002.

145. K. Herzer, T. M. Ganten, H. Schulze-Bergkamen, A. Grosse-Wilde, R. Koschny, P. H. Krammer, and H. Walczak, “Transforming growth factor  can mediate apoptosis via the expression of trail in human hepatoma cells,” *Hepatology*, vol. 42, no. 1, pp. 183–192.

151. M. Swiatkowska, J. Szemraj, and C. S. Cierniewski, “Induction of pai-1 expression by tumor necrosis factor  in endothelial cells is mediated by its responsive element located in the 4g/5g site,” *The FEBS Journal*, vol. 272, no. 22, pp. 5821–5831.

159. N. Kanda, T. Shimizu, Y. Tada, and S. Watanabe, "Il-18 enhances ifn--induced production of cxcl9, cxcl10, and cxcl11 in human keratinocytes," *European Journal of Immunology*, vol. 37, no. 2, pp. 338–350.
